# Study of microbe-microbe interactions between the sexually transmitted parasite *Trichomonas vaginalis* with the cervicovaginal bacteria *Lactobacillus iners*

**DOI:** 10.64898/2026.07.14.738301

**Authors:** Samantha Smedshammer, Bryn Baxter, Geraldine J. Briceno, Kathryn E. Morales, Anel Lizcano, Ty’Tianna Clark, Dustin Willard, Angelica M. Riestra

## Abstract

*Trichomonas vaginalis* is the leading cause of non-viral sexually transmitted infections and it is associated with comorbidities that affect female health. *Lactobacillus iners* is one of the most predominant bacteria in the cervicovaginal microbiome. As so, both microbes are likely to encounter one another upon *T. vaginalis* infection. To our knowledge, the interaction of both microbes has not been previously investigated. Here, we report that *T. vaginalis* and *L. iners* bind to one another at early time points of co-incubation. Using imaging flow cytometry and scanning electron microscopy, we capture the dynamics of this microbe-microbe association. We observed active remodeling of the *T. vaginalis* cell surface leading to thin-membrane protrusions that make contact with *L. iners*. Larger *T. vaginalis* membrane extensions that surround and engulf *L. iners* were also visible. These *T. vaginalis*-*L. iners* interactions ultimately lead to a reduction of *L. iners* viability while *T. vaginalis* viability was unaffected by exposure to *L. iners*. Inhibition of actin polymerization blocked *T. vaginalis* antibacterial activity against *L. iners*. Together our findings reveal novel insight about *T. vaginalis*-*L. iners* interactions and highlight a new *T. vaginalis* pathogenic effect.

## Introduction

The parasite *Trichomonas vaginalis* causes trichomoniasis, a neglected parasitic infection of the U.S. This designation arises as result of *T. vaginalis* being understudied and its disproportionate affliction of racial and ethnic minorities [1]. It is estimated that 2.6 million people are currently infected with *T. vaginalis* in the U.S. [2] and that there are 156 million new cases each year worldwide [3], making *T. vaginalis* the most common cause of non-viral sexually transmitted infections [3]. Women have a 5.2-fold increased risk of *T. vaginalis* infection compared to men [4]. The higher prevalence among women is concerning as many comorbidities are also experienced by them. For example, *T. vaginalis* infection increases the risk of preterm birth, giving birth to a low-birth-weight infant [5, 6], premature rupture of the membranes [6], increased risk of cervical cancer [7, 8] and HIV acquisition [9]. *T. vaginalis* is also associated with bacterial vaginosis, a condition characterized by an imbalance in the vaginal microbiota [10].

Women harbor one of five bacterial community groups, known as community state types (CSTs) I-V [11]. CSTs I-III are dominated by a *Lactobacillus* species whereas CST-IV has reduced amounts of *Lactobacillus* species and increased amounts of anaerobic bacteria such as *Gardnerella vaginalis* and *Prevotella bivia* [11]. A CST-IV is also characteristic of women with bacterial vaginosis [11, 12]. Interestingly, many of the gynecologic and obstetric sequalae observed upon *T. vaginalis* infection-including preterm birth and increased HIV infection are also associated with bacterial vaginosis [13–17]. Furthermore, *T. vaginalis* is increasingly detected in women with clinically diagnosed bacterial vaginosis [18, 19] and with cervicovaginal bacteria characteristic of bacterial vaginosis [20, 21]. It is largely unknown if these associations exist as a result of 1) *T. vaginalis* infection directly driving a reduction in the abundance of *Lactobacillus* species in the cervicovaginal microbiome which in turn facilitates the outgrowth of CST-IV/bacterial vaginosis bacteria and/or 2) whether modification of the cervicovaginal microbiome after *T. vaginalis* colonization synergizes in causing comorbidities associated with trichomoniasis. Clinically, reduced amounts of lactobacilli in women with trichomoniasis has also been reported [22]. *In vitro T. vaginalis* has been found to reduce the viability of some *Lactobacilli* species [23, 24]. However, to our knowledge the interaction of *T. vaginalis* with *Lactobacillus iners* has never been characterized. Additionally, the experimental conditions shown in this study differ from prior investigations of *T. vaginalis*-lactobacilli interactions [23, 24].

The *T. vaginalis*-*L. iners* microbe-microbe interaction is of high significance to study given that *L. iners* is potentially the most prevalent vaginal bacteria in women of reproductive age [25–27]. *L. iners* is also the most common cervicovaginal bacteria in Black women [11, 27, 28] and Hispanic women in the U.S. [11, 29, 30] and in pregnant women of either African or European ancestry [29]. While *L. crispatus* is thought to play a protective role in female health given that its presence is associated with decreased risk of viral, fungal, and bacterial infections, adverse pregnancy outcomes, and cervical cancer [14, 27], less is known about the contribution of *Lactobacillus iners* towards female reproductive health [26, 31]. *L. iners* has been understudied largely due to difficulty in culturing the bacteria. However, a recent technical advancement has improved bacterial growth in liquid media [32]. In this work, we build on this knowledge and report an improvement to quantification of *L. iners* on agar plates. Our study also generates increased knowledge about the functional properties of *L. iners* by investigating its effects against a protozoan pathogen, as little is also known about the effect of lactobacilli against *T. vaginalis*.

Out of all the CSTs, a CST-III dominated by *L. iners* is the most unstable, with documented shifts in women towards a CST-IV/bacterial vaginosis-like community [33, 34]. Thus, perturbations to *L. iners* may have magnified consequences in women with cervicovaginal microbiomes dominated by this bacteria. Furthermore, out of all the cervicovaginal lactobacilli, *L. iners* can also be detected in cervicovaginal microbiomes dominated by other lactobacilli [11, 35, 36], but it is largely unknown how fluctuations of *L. iners* amounts affect these women. In this study, we tested if *T. vaginalis* directly binds to *L. iners* and observed the nature of these interactions using imaging flow cytometry and scanning electron microscopy. In conditions that maximize the growth of both microbes, we also tested how *T. vaginalis*-*L. iners* co-incubation affect the viability of both microorganisms. We found that *T. vaginalis* potently and rapidly kills *L. iners*, whereas *L. iners* exposure did not affect the viability of *T. vaginalis*. We further investigate the mechanisms contributing to *T. vaginalis* antibacterial activity against *L. iners*.

## Materials and Methods

### Optimization of *L. iners* growth

To establish the optimal growth conditions for *Lactobacillus iners*, various agar types and supplements were tested. The base media included De Man-Rogosa-Sharpe (MRS) agar (BD Difco, Catalog #: 288210), Columbia blood agar base (Thermo Fisher Scientific, Catalog #: OXCM0233B, and NYC-III agar base (Himedia, Catalog #: M1348). All agar types were prepared in Milli-Q water, sterilized by autoclaving, and cooled to approximately 50°C prior to the addition of supplements: 10% (v/v heat-inactivated horse serum (Thermo Fisher Scientific, Catalog #: NC1723618), 5% (v/v) defibrinated horse blood (Thermo Fisher Scientific, Catalog #: R54012), 1.1 mM L-glutamine (Thermo Fisher Scientific, Catalog #: 25030081), and 4 mM L-cysteine (Sigma-Aldrich, Catalog #: 501790329). After supplement addition, agar was poured into sterile 100 mm petri dishes (approximately 10 mL per plate) and allowed to solidify at room temperature for two days. Serial ten-fold dilutions of *L. iners* (ATCC 55195) were performed and plated onto the different types of agar plates. Afterwards, they were incubated anaerobically at 37°C using BD GasPak™ EZ anaerobe containers (BD, Catalog #: B260001) and observed for colony formation.

### Imaging Flow Cytometry

*Trichomonas vaginalis* (strain MSA 1132; Mt. Dora, FL, USA) was cultured in Diamond’s medium and *Lactobacillus iners* (ATCC 55195) was cultured in De Man-Rogosa-Sharpe (MRS) broth (RPI, Catalog #: L11000) supplemented with 4 mM L-cysteine and 1.1 mM L-glutamine (MRS-CQ). On the day prior to co-incubation, *T. vaginalis* was cultured in Diamond’s media without the routine addition of penicillin G and streptomycin (Diamond’s media −P/−S). On the day of the experiment, both organisms were resuspended in Live Cell Imaging Solution (Thermo Fisher Scientific, Catalog #: A59688DJ) and both microbes were fluorescently pre-labeled using Biotium CellBrite® Fix 488 (green fluorescent; Biotium Catalog #: 501964510) for *L. iners* and Biotium CellBrite® Fix 640 (red fluorescent; Biotium Catalog #: 30089T) for *T. vaginalis*. Co-incubation was performed by combining ~2×10^7^ cells of *L. iners* (resuspended in 250 µL) with 1×10^6^ *T. vaginalis* (resuspended in 250 µL) for a target 20:1 multiplicity of infection (MOI), in a final volume of 500 µL. Samples were incubated at 37°C under anaerobic conditions using BD GasPak™ EZ anaerobe containers (BD, Catalog #: B260001) for 30 minutes and 1 hour. Stained control samples and unstained control samples containing a single type of microbe were incubated in parallel to serve as fluorescence positive and negative controls, respectively, for gate setting. At each time point, samples were fixed by resuspension in 250 µL of 4% formaldehyde (Electron Microscopy Sciences, Catalog #: 15710) diluted in FACS buffer (PBS with 5% FBS, 10 mM EDTA, and 0.1% sodium azide) and incubated for 15 minutes on ice. Fixed samples were then washed three times with FACS buffer and resuspended in a final volume of 150 µL. ImageStream (Amnis) was utilized to acquire 5,000 events from each sample. Data was analyzed using IDEAS® 6.3 software with the Internalization Wizard and AI-based analysis tools to differentiate cell populations based on hundreds of measured features such as size, shape, texture, and fluorescence intensity.

### Scanning electron microscopy

~2×10^7^ *L. iners* CFUs and 1×10^6^ *T. vaginalis* MSA 1132 cells were mixed to achieve a target MOI of 20:1 on top of poly-L-lysine-coated glass coverslips (BD, Catalog #: 08774384) (deposited within 24-well plates) at 37°C for 30 minutes, 1 hour, and 1.5 hours under anaerobic conditions. Afterwards, coverslips were fixed in 2.5% glutaraldehyde prepared in 0.1 M cacodylate buffer (pH 7.2) for 30 minutes at room temperature. After fixation, samples were washed three times in 0.1 M cacodylate buffer and post-fixed in 1% osmium tetroxide for 30 minutes. Samples were then dehydrated using a graded ethanol series: 50%, 75%, 95% (twice), and 100% (twice), with each step lasting 15 minutes. Dehydrated samples were processed using a critical point dryer, mounted on aluminum stubs with double-sided carbon adhesive tape, and sputter-coated with a 6 nm layer of platinum. Imaging was conducted using a Quanta 450 FEG scanning electron microscope operating at 5 kV at the SDSU Electron Microscopy Facility. Image annotation was performed using Adobe Photoshop 2025. Image analysis and quantification of bacterial morphology were performed using FIJI (ImageJ) software.

### Total killing assays

To evaluate the viability of *L. iners* and *T. vaginalis* upon co-incubation, we aimed to co-incubate both microbes at an MOI of 20 *L. iners* to 1 *T. vaginalis* by aliquoting ~2×10^7^ *L. iners* CFUs and 1×10^6^ *T. vaginalis* (MSA 1132) in a final volume of 0.5 mL. To achieve this a master mix of each microbe was resuspended in their optimal microbiological media and 250 µL of each microbe stock was aliquoted in a well of a 24-well plate to generate a final 1:1 mixture of MRS+QC media (optimal *L. iners* microbiological media) and Diamond’s media (optimal *T. vaginalis* microbiological media), which we refer to as 50/50 media. A 3 hour co-incubation at 37°C was then performed under anaerobic conditions maintained using the BD GasPak™ EZ Anaerobe Container System. As single-microbe controls, each microorganism was incubated alone in its optimal growth medium and in the 50/50 media. The latter served as the “*L. iners* only” control against which growth was compared by normalization to “100%”. At the end of the incubation, samples were transferred to sterile 1.5 mL microcentrifuge tubes. 500 µL of Accutase (Innovative Cell Technologies, catalog #: NC9839010) was then added to each well to recover any remaining adherent cells and incubated for 30 minutes at 37°C/5%CO₂. Wells were then washed by pipetting up and down around the well 10 times to harvest remaining contents and pooled with the sample in the microcentrifuge tubes. Afterwards, 100 µL of heat-inactivated horse serum was added to these tubes to neutralize Accutase activity. Tubes were then vortexed for 5 seconds. *L. iners* viability was determined by performing serial ten-fold dilutions in PBS and plating on Columbia Blood Agar supplemented with 5% sheep blood, 4 mM L-cysteine, and 1.1 mM L-glutamine (CBA+CQ). CBA+CQ plates were incubated anaerobically at 37°C for 48 hours to assess formation of colony forming units (CFUs). *T. vaginalis* viability was quantified via performing hemocytometer cell counts.

### Cytochalasin D Inhibitor Assay

Total killing assays were performed in 24-well plates as described above in the presence of the actin polymerization inhibitor cytochalasin D (Sigma Aldrich, catalog #: C8273) or DMSO vehicle control. Cytochalasin D or DMSO were added to the *T. vaginalis* or Diamonds media master mix to achieve a final concentration of 60 µM in a final volume of 0.510 ml upon mixing 0.260 ml of the *T. vaginalis* or Diamonds media master mix with 0.250 ml of *L. iners* or MRS+CQ media. Co-incubations were similarly performed by combining ~2×10^7^ *L. iners* CFUs with 1×10^6^ *T. vaginalis* (MSA 1132). After a 3 hour incubation, each microbe’s viability was assessed as described above.

### Data Analysis

GraphPad Prism (Version 10) was used for data visualization and statistical analysis.

## Results

### T. vaginalis associates with L. iners

To characterize the early microbe-microbe events of *T. vaginalis* with *L. iners,* we used ImageStream, an imaging flow cytometer, that allowed us to take single cell images and quantify cellular observations. *T. vaginalis* and *L. iners* were fluorescently prelabeled with Biotium CellBrite Fix 488 (green) and CellBrite Fix 640 (red), respectively, and then co-incubated under anaerobic conditions for 30 minutes and 1 hour. Each microbe was also singly incubated in parallel as microbe only controls. ImageStream was then utilized to acquire single cell images, and IDEAS software was employed to quantify the imaging results. First, the *L. iners* and *T. vaginalis* “cells” populations were identified by gating in-focus cells based on the Gradient RMS feature on the IDEAS software. Next, the two populations were further defined based on the Area morphology feature. Afterwards, given that both microbe populations were stained with two different fluorescent dyes, the unstained and singly stained single microbe controls were used to define the positively stained fluorescent populations for each microbe using the IDEAS Intensity feature. Finally, the occurrence of double positive events indicative of red fluorescent *T. vaginalis* interaction with green fluorescent *L. iners* was quantified. A statistically significant increase in the amount of double positive counts was detected from 30 minutes to 1 hour of co-incubation leading to a 3-fold increase in *T. vaginalis*-*L. iners* association events at the later time point (**Fig. 1A**). Furthermore, observation of these images revealed co-localization between *T. vaginalis* and *L. iners* by 30 minutes of co-incubation and potential internalization of *L. iners* by *T. vaginalis* at 1 hour (**Fig. 1B**).

**Figure 1.**
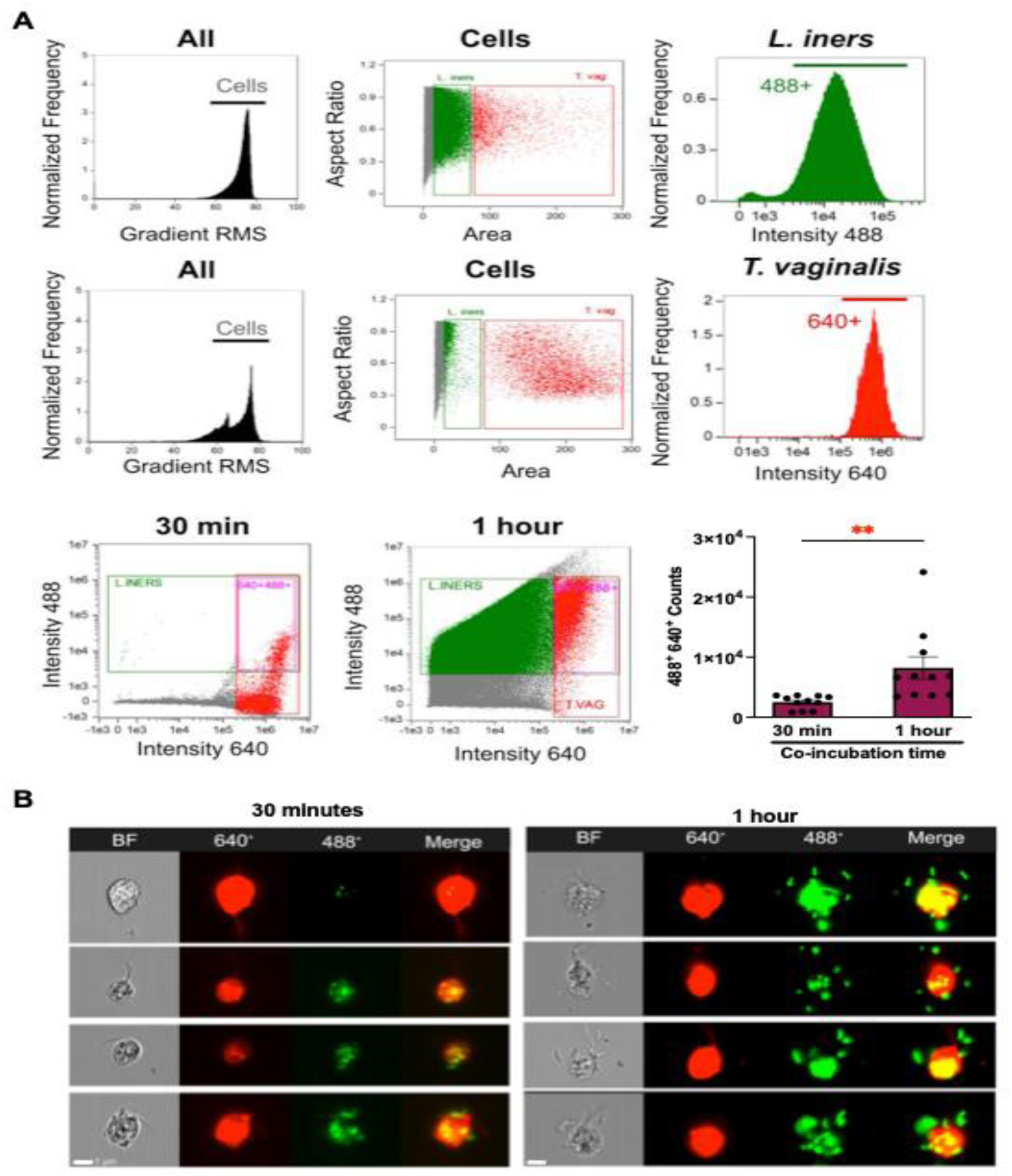
*T. vaginalis* interacts with *L. iners*. *L. iners* was exposed to *T. vaginalis* at MOIs of 3-16 for 30 minutes or 1 hour and microbe-microbe interactions were quantified by imaging flow cytometry. (**A**) Gating strategy to identify green fluorescent *L. iners* (488^+^), red fluorescent *T. vaginalis* (640^+^), and red fluorescent *T. vaginalis* associated with green fluorescent *L. iners* (double positive 488^+^ 640^+^). In-focus cells were first selected (gradient RMS), followed by differential morphology based on area, and then fluorescence signal above background controls (488^+^ or 640^+^). Graph shows quantification of the double positive population. Bars show the pooled mean of 3 biologically independent experiments with standard error of the mean displayed as error bars. Individual values from 4 technical replicates performed in each biologically independent experiment are shown as dots. **=p-value <0.01. (**B**) Representative ImageStream images of the double positive population quantified in A. Individual brightfield (BF), red fluorescent channel (640^+^), green fluorescent channel (488^+^), and a merge of both fluorescence channels is shown. Red fluorescent *T. vaginalis* in contact with green fluorescent *L. iners* after 30 minutes and 1 hr co-incubation can be observed, with yellow co-localization in the merge visible upon *T. vaginalis*-*L. iners* association.

### Dynamics of *T. vaginalis-L. iners* interactions

To further examine *T. vaginalis*-*L. iners* interactions at higher resolution, we employed scanning electron microscopy (SEM). Co-incubation of *T. vaginalis* with *L. iners* led to visible progressive membrane remodeling events on the parasite cell surface. Striking *T. vaginalis* membrane protrusions that formed lattice-like structures around *L. iners* were detected as early as 30 minutes of co-incubation (**Fig. 2A-2B**). The thin membrane protrusions could also be observed extending outward from the *T. vaginalis* surface towards *L. iners* (**Fig. 2C-2D**). These images highlight early recognition and physical tethering of *L. iners* by *T. vaginalis*. We also observed larger regions of the *T. vaginalis* membrane wrapping around *L. iners* (**Fig. 2E-2F**). As early as 30 minutes of co-incubation, we also visualized whole *L. iners* being internalized by *T. vaginalis* (**Fig. 3**). *L. iners* was observed located over a membrane indentation on the *T. vaginalis* surface with membrane remodeling visible beneath the bacteria (**Fig. 2A**). *L. iners* could also be found inside similar *T. vaginalis* membrane indentations with signs of membrane ruffling at the outer diameter (**Fig. 3B 3C** and **Fig. 3D-3E**).

**Figure 2.**
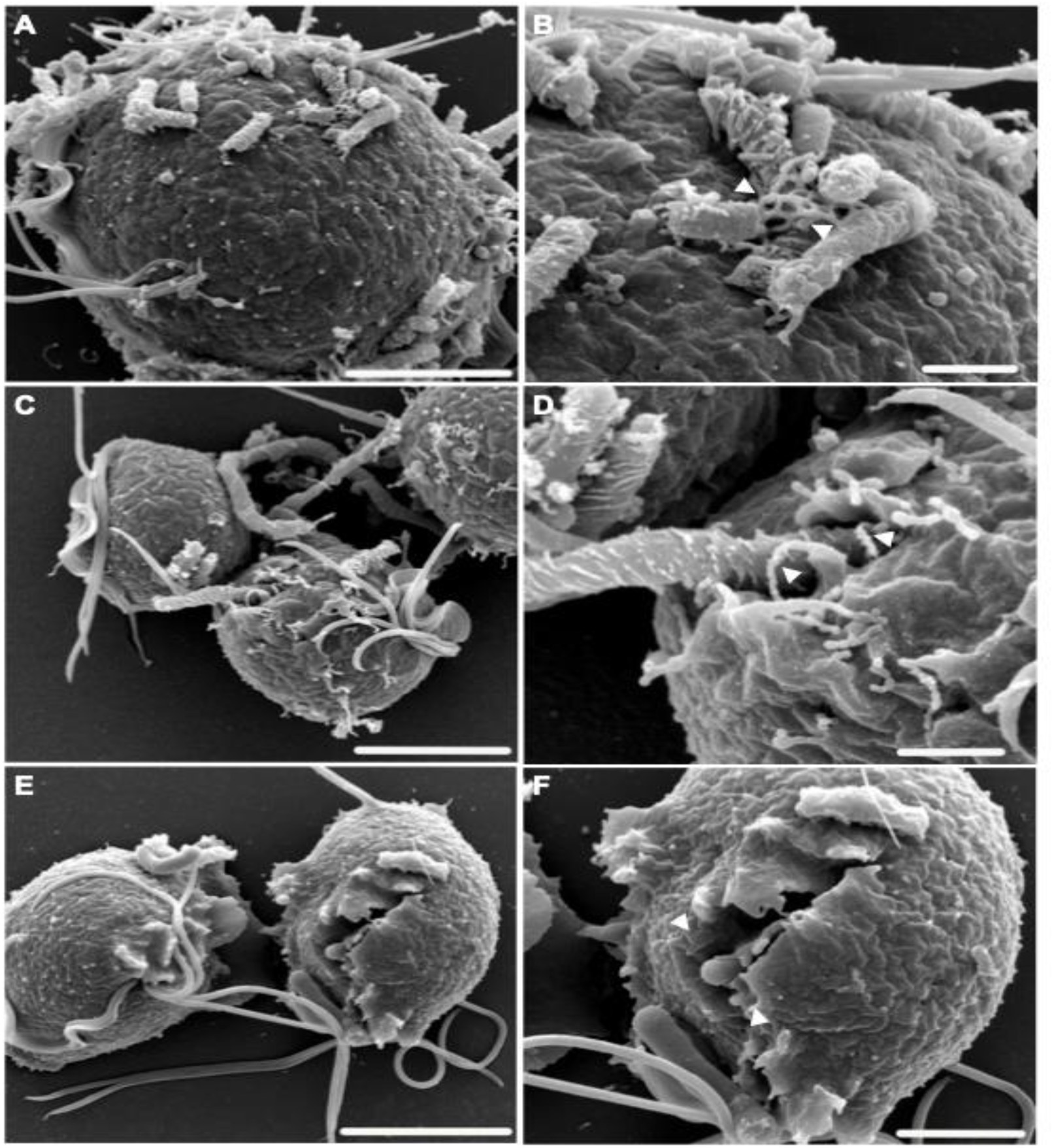
*T. vaginalis* membrane reorganizes around *L. iners*. *L. iners* was co-incubated with *T. vaginalis* on poly-L lysine coated coverslips at MOIs of 4 to 32 under anaerobic conditions for 30 minutes, 1hour, and 1.5 hours. Samples were fixed and imaged by scanning electron microscopy. Pairs of an image and a zoom-in image next to it are shown. SEM micrographs reveal progressive membrane remodeling events on *T. vaginalis* cell surface during co-incubation with *L. iners* as indicated by the white arrows in the zoom-in image. Note observation of (**A-B**) lattice-like membrane structures wrapping around *L. iners*, (**B-C**) membrane protrusions extending towards *L. iners*, and (**C**) membrane engulfment of *L. iners.* Scale bars: (**A-B** and **C-D**) 5µm, 2µm and (**E-F**) 5µm, 1 µm.

**Figure 3.**
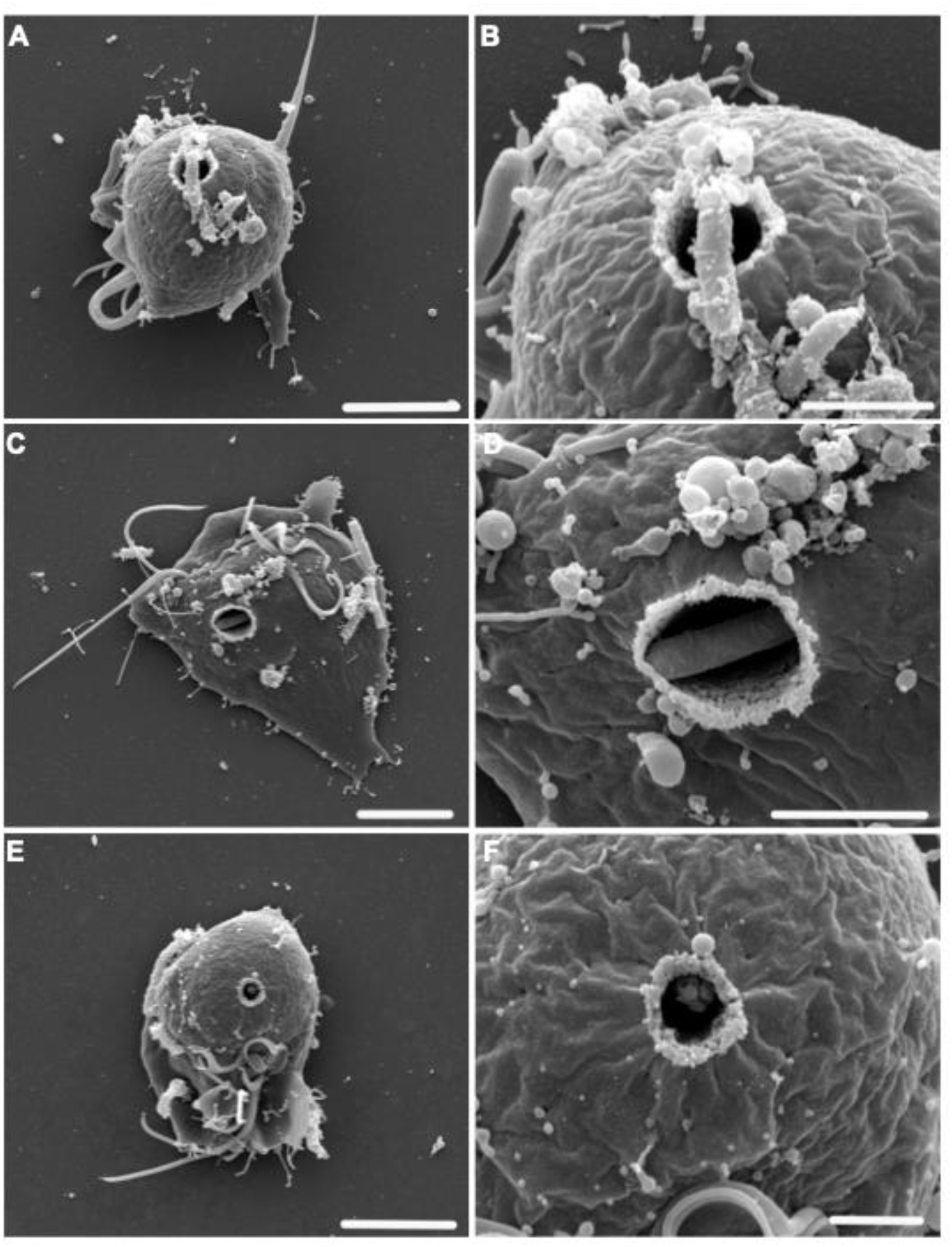
*L. iners* is internalized by *T. vaginalis*. *L. iners* was co-incubated with *T. vaginalis* on poly-L lysine coated coverslips at MOIs of 4 to 32 under anaerobic conditions for 30 min 1hr and 1.5 hours. Samples were fixed and imaged by scanning electron microscopy. Pairs of an image and a zoom-in image next to it are shown. SEM micrographs of co-incubation conditions show *L. iners* being internalized by *T. vaginalis* through membrane indentations on the parasite surface. Scale bars: (A-B and C-D) 5µm, 2µm (E-F) 5µm, 1µm.

### *T. vaginalis* alters *L. iners* surface morphology

To characterize the outcome of these interactions, we performed quantitative assessment of the bacterial morphology in SEM micrographs (**Fig. 4**). In control conditions where *L. iners* was incubated alone, bacterial cells displayed smooth, intact rod-like structures with well-defined surface topology at both 30 minutes and 1 hour (**Fig. 4A**). In contrast, upon *L. iners* co-incubation with *T. vaginalis*, a distinct “zebra-like” striated surface pattern was observed on the *L. iners* surface at both 30 minutes and 1 hour of exposure to the parasite (**Fig. 4A**). This “zebra-like” striated appearance was detected at very low levels in single-culture controls and was reproducibly higher upon exposure of *L. iners to T. vaginalis* (**Fig. 4B**). These analyses suggested that interaction with *T. vaginalis* may induce structural damage to *L. iners* and that this effect is detected early-on in these microbe-microbe interactions.

**Figure 4.**
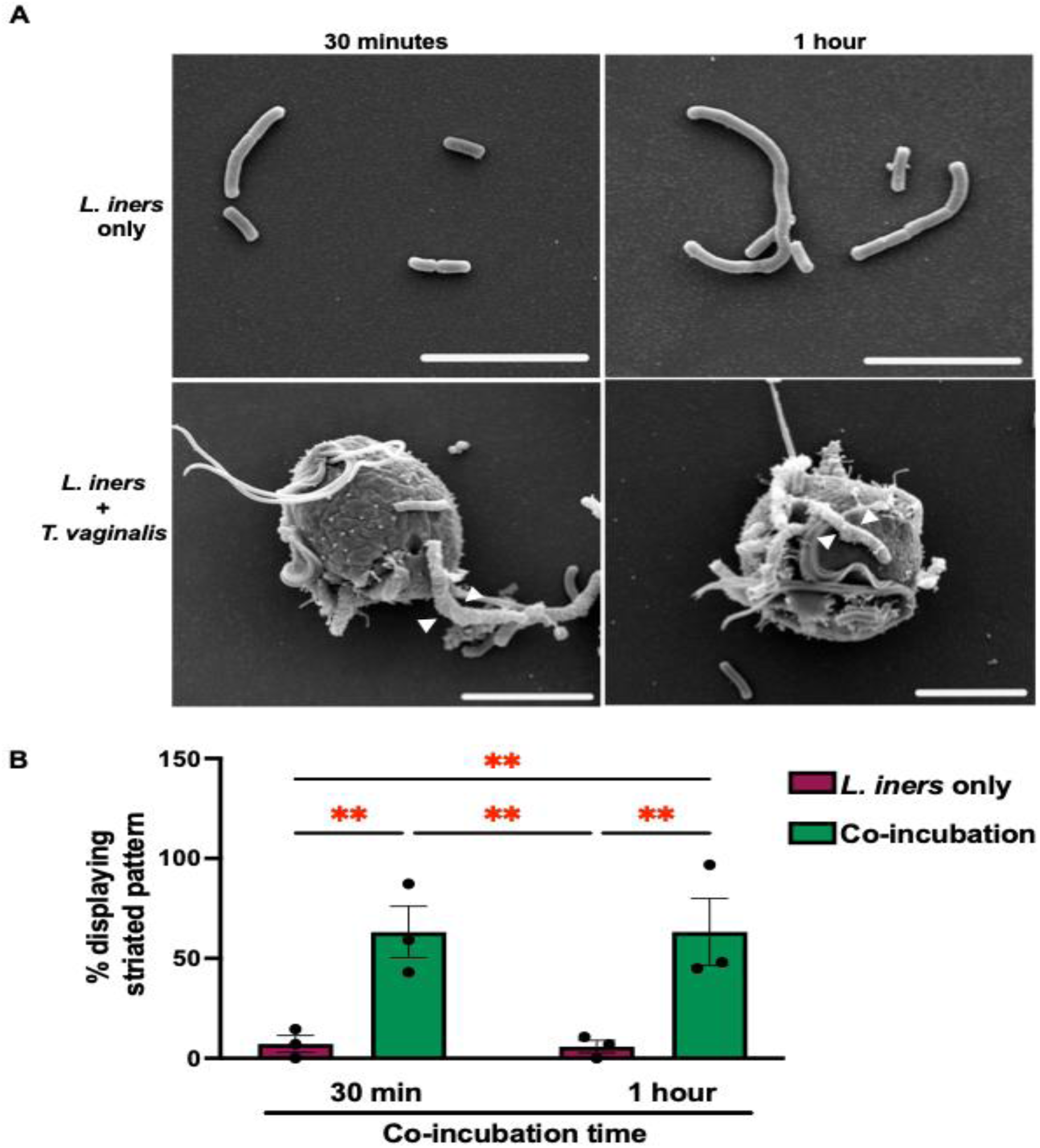
*T. vaginalis* alters the membrane morphology of *L. iners*. *L. iners* was co-incubated with *T. vaginalis* on poly-L lysine coated coverslips at MOIs of 4 to 32 under anaerobic conditions for 30 minutes and 1 hour. (**A**) Representative SEM micrographs of *L. iners* only (top panel) and co-incubation conditions (bottom panel) at 30 minutes (left) and 1 hour (right). Co-incubation of *L. iners* with *T. vaginalis* led to the increased detection of a striated pattern on the cell surface of *L. iners* (white arrows). (**B**) The percent of *L. iners* that displayed the striated cell surface pattern was quantified. Graph shows pooled mean of 3 biologically independent experiments at 30 minutes and 1 hour. A total of 77 individual bacteria per time point were quantified. Standard error of the mean is displayed as error bars. Individual values are shown as dots. ** = p < 0.001.

### Close contact with *T. vaginalis* is required to induce *L. iners* surface remodeling

To determine whether physical interaction between *T. vaginalis* and *L. iners* was required to inflict the *L. iners* surface alterations, we employed Cerillo’s Co-Culture Duet systems. This system contains two chambers separated by a 0.2 μm membrane filter that permits diffusion of soluble molecules while preventing direct cell-cell contact (**Fig. 5A**). Each microbe was incubated alone, co-incubated together, or cultured in the adjacent chambers separated by the membrane (**Fig. 5B**). In the single microbe controls and the co-incubation condition, media was added to the adjacent chamber to maintain the same final volume across both chambers. Following incubation, each treatment condition was processed and imaged using SEM, and the percentage of *L. iners* displaying the characteristic striated surface morphology was quantified. Direct co-incubation of *L. iners* with *T. vaginalis* resulted in a statistically significant increase in the proportion of bacteria displaying the striated phenotype compared to *L. iners* cultured alone (**Fig. 5C**). In contrast, physical separation of the two microorganisms eliminated this phenotype, with *L. iners* exhibiting surface striations at similar levels comparable to the bacteria only condition. These findings demonstrate that induction of the altered *L. iners* surface morphology requires contact with *T. vaginalis*.

**Figure 5.**
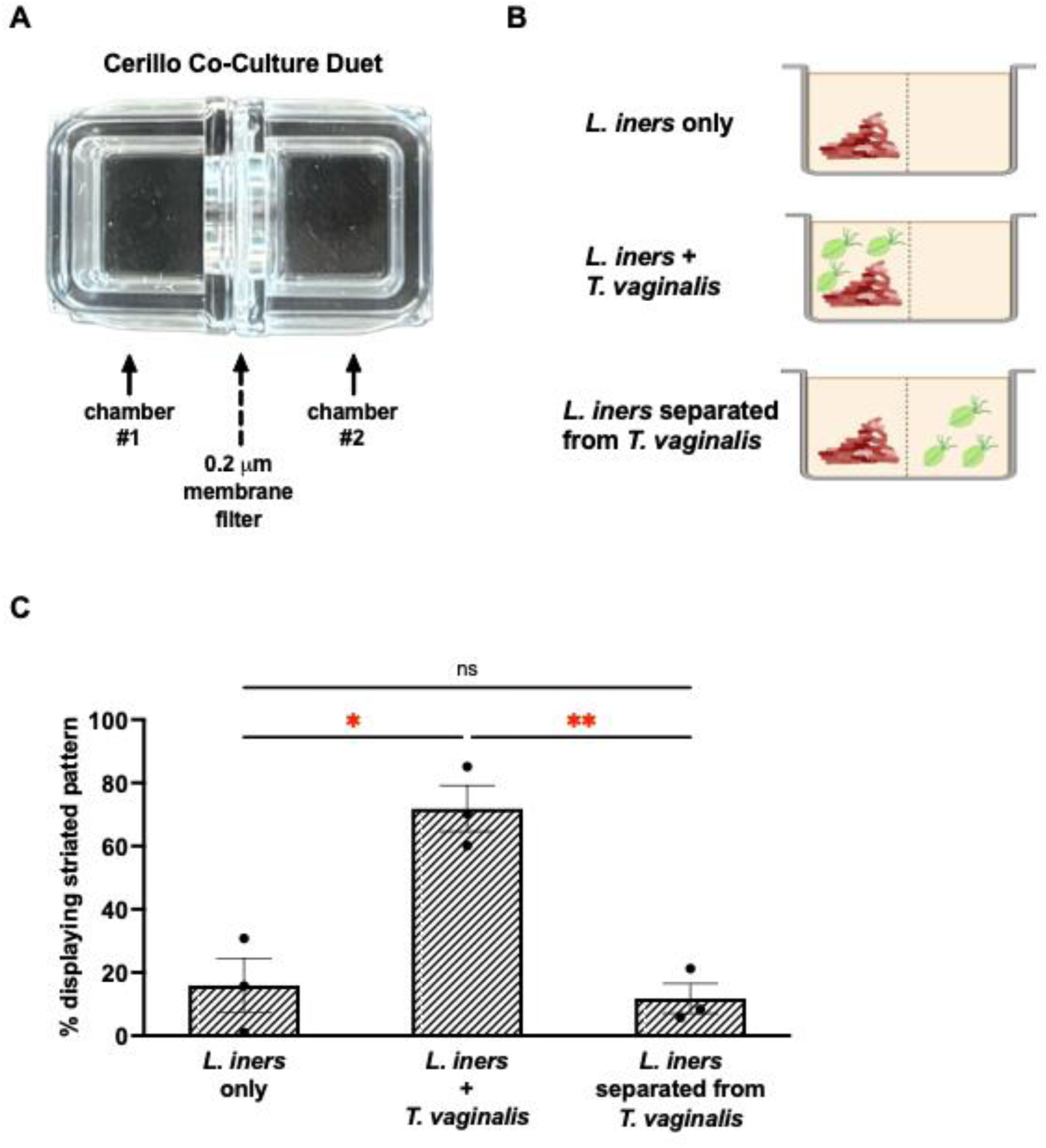
Physical separation reduces *L. iners* membrane striation morphology. *L. iners* was co-incubated with *T. vaginalis* under varying conditions in the Cerillo’s Duet Co-Culture System at MOIs of 12 to 57 under anaerobic conditions for 1.5 hours. The percent of *L. iners* that displayed a striated pattern on the cell surface was quantified. (**A**) Shows an image of the Cerillo Co-Culture Duet system which contains two chambers separated by a membrane filter. (**B**) Shows a schematic of the experimental conditions tested to perform the quantification of *L. iners* striations shown in C. (**C**) After co-incubation in the conditions shown in B, the microbes were transferred onto poly-L lysine coated coverslips, were fixed, and imaged using SEM. SEM micrographs were analyzed to quantify the percent of *L. iners* with a striated surface morphology (as in Fig. 4). Graph shows pooled mean of 3 biologically independent experiments. A total of 508 individual bacteria per condition were quantified. Standard error of the mean is displayed as error bars. Individual values are shown as dots. *=p-value<0.1 **=p-value<0.01.

### *T. vaginalis* exhibits selective antibacterial activity against cervicovaginal bacteria

To determine whether the interactions observed between *T. vaginalis* and *L. iners* resulted in altered microbial viability, we quantified the recovery of both microorganisms after 3 hours of co-incubation under anaerobic conditions at MOIs 26-28 (**Fig. 6**). Recovery of *L. iners,* measured by formation of colony forming units (CFUs) on Columbia Blood Agar plates supplemented with L-cysteine and L-glutamine (CBA +CQ), was dramatically reduced following co-incubation with *T. vaginalis* leading to approximately 95% fewer CFUs recovered compared to *L. iners* cultured alone (**Fig. 6A**). In contrast, recovery of *T. vaginalis*, measured via hemocytometer cell counts, was not altered by the presence of *L. iners*, leading to similar percent recovery of *T. vaginalis* compared to the *T. vaginalis* only control (**Fig. 6B**). Together, these results indicate that while *T. vaginalis* substantially reduced *L. iners* viability, its own survival remained unaffected by exposure to *L. iners*.

**Figure 6.**
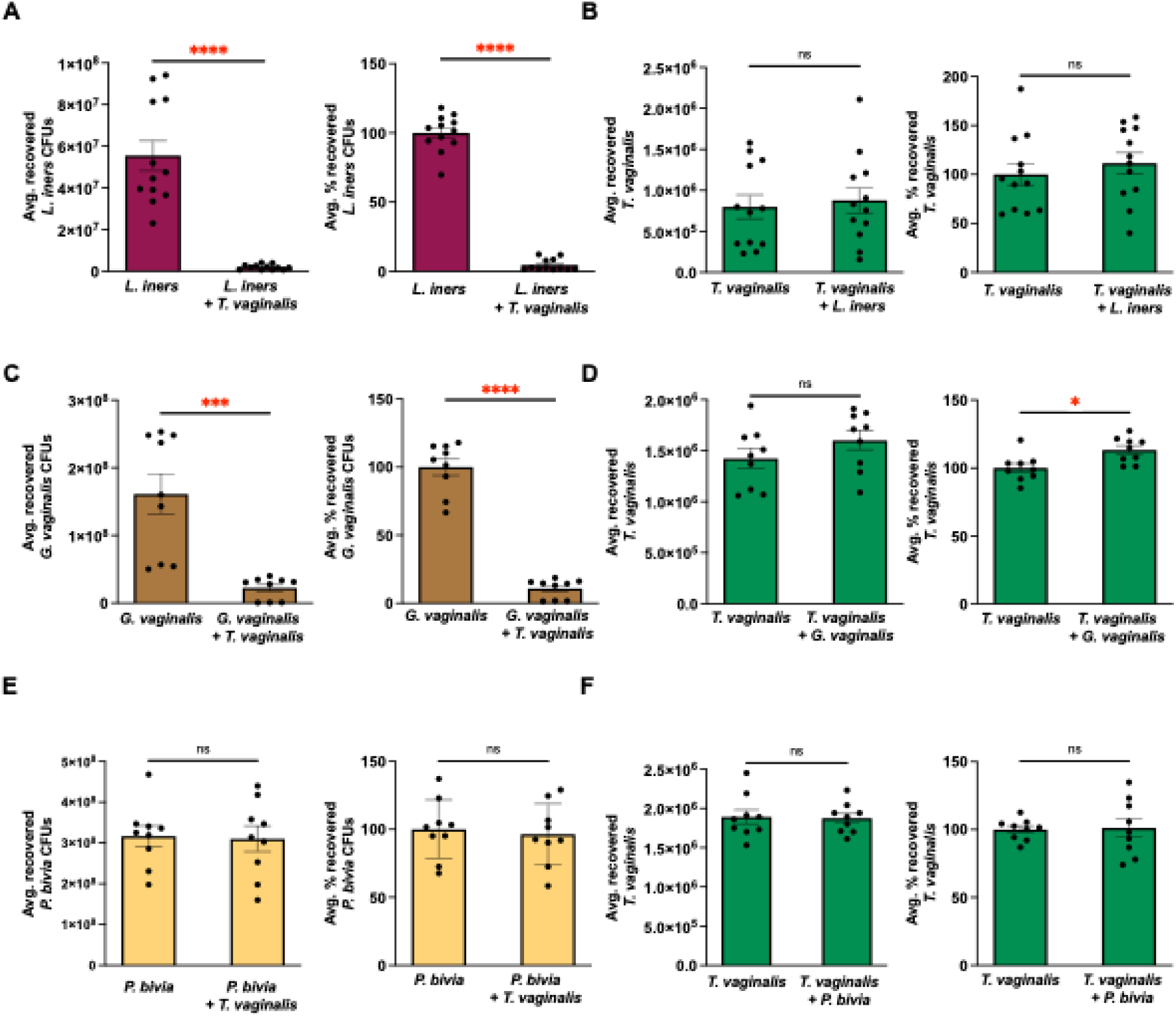
*T. vaginalis* exhibits antibacterial activity against cervicovaginal bacteria. *T. vaginalis* was co-incubated with *L. iners* (A-B), *G. vaginalis* (C-D), or *P. bivia* (E-F) for 3 hours under anaerobic conditions at MOIs of 26–28 *L. iners*, 15–31 *G. vaginalis,* and 23–32 *P. bivia* to *T. vaginalis*. As controls, each microbe was incubated alone in the same 50/50 experimental medium consisting of equal volumes of the optimal growth medium for each microbe. (**A, C, E**) Bacterial viability was quantified by assessing formation of colony forming units (CFUs) on Columbia Blood agar +QC for *L. iners*, Casman’s Medium Base agar for *G. vaginalis*, and Tryptic Soy agar for *P. bivia* (left graph) and normalized to the corresponding bacteria-only control (right graph). (**B, D, F**) *T. vaginalis* viability was quantified by hemocytometer counts (left graph) and normalized to the corresponding *T. vaginalis*-only control (right graph). Graphs show pooled data from three biologically independent experiments, each performed with three technical replicates. Error bars show the standard error of the mean, and individual technical replicate values are shown as dots. ****=p-value < 0.0001, ***=p-value < 0.001,*=p-value < 0.05; ns, not significant (p-value > 0.05). Co-incubation with *T. vaginalis* reduced the viability of *L. iners* and *G. vaginalis*, whereas *P. bivia* viability was unaffected. *T. vaginalis* viability was not inhibited by co-incubation with any bacterial species.

To determine whether this antibacterial activity extended to other members of the vaginal microbiota, we similarly performed total killing assays with *Gardnerella vaginalis* and *Prevotella bivia* at MOIs 15–31 and 23–32, respectively under 50/50 media conditions. Co-incubation with *T. vaginalis* resulted in a significant reduction in *G. vaginalis* viability, with an approximately 89% decrease in recovered CFUs compared to *G. vaginalis* cultured alone (**Fig. 6C**). *T. vaginalis* exhibited a slight statistically significant increase in total cell counts with a 119% percent recovery following co-incubation with *G. vaginalis* compared to the *T. vaginalis* only condition (**Fig. 6D**). In contrast, *T. vaginalis* co-incubation with *P. bivia* had no significant effect on either bacterial CFU recovery (**Fig. 6E**) or *T. vaginalis* cell recovery (**Fig. 6F**). Together, these findings demonstrate that *T. vaginalis* possesses selective antibacterial activity, efficiently reducing the viability of *L. iners* and *G. vaginalis* while not affecting *P. bivia*, and that the presence of these bacteria do not compromise parasite viability.

### Actin polymerization contributes to *T. vaginalis* killing of *L. iners*

Given that we had observed active remodeling of the *T. vaginalis* membrane upon exposure to *L. iners* (**Fig. 2**) and bacterial uptake (**Fig. 2E-2F** and **Fig. 3**), coupled with antibacterial activity displayed against *L. iners* (**Fig. 6A**), we predicted that actin dynamics contributed to *T. vaginalis* antibacterial effect against *L. iners*. To investigate this, *T. vaginalis* was treated with the actin polymerization inhibitor cytochalasin D upon co-incubation with *L. iners*. Similar to the antibacterial effects observed in **Fig. 6A**, *T. vaginalis*-*L. iners* co-incubation in the presence of vehicle control led to a statistically significant 95% reduction in *L. iners* CFUs compared to *L. iners* treated with the vehicle control (**Fig. 7A**) whereas inhibition of parasite actin polymerization led to a protective effect restoring *L. iners* recovery to the amounts observed in the *L. iners* only control (**Fig. 7A-7B**). Importantly, cytochalasin D treatment had no significant effect on *T. vaginalis* viability leading to similar amounts of parasite recovered compared to the *T. vaginalis*-*L. iners* co-incubation condition (**Fig. 7C-7D**), indicating that the rescue of *L. iners* was not due to loss of parasite viability. These findings demonstrate that *T. vaginalis* actin polymerization is required for efficient killing of *L. iners*.

**Figure 7.**
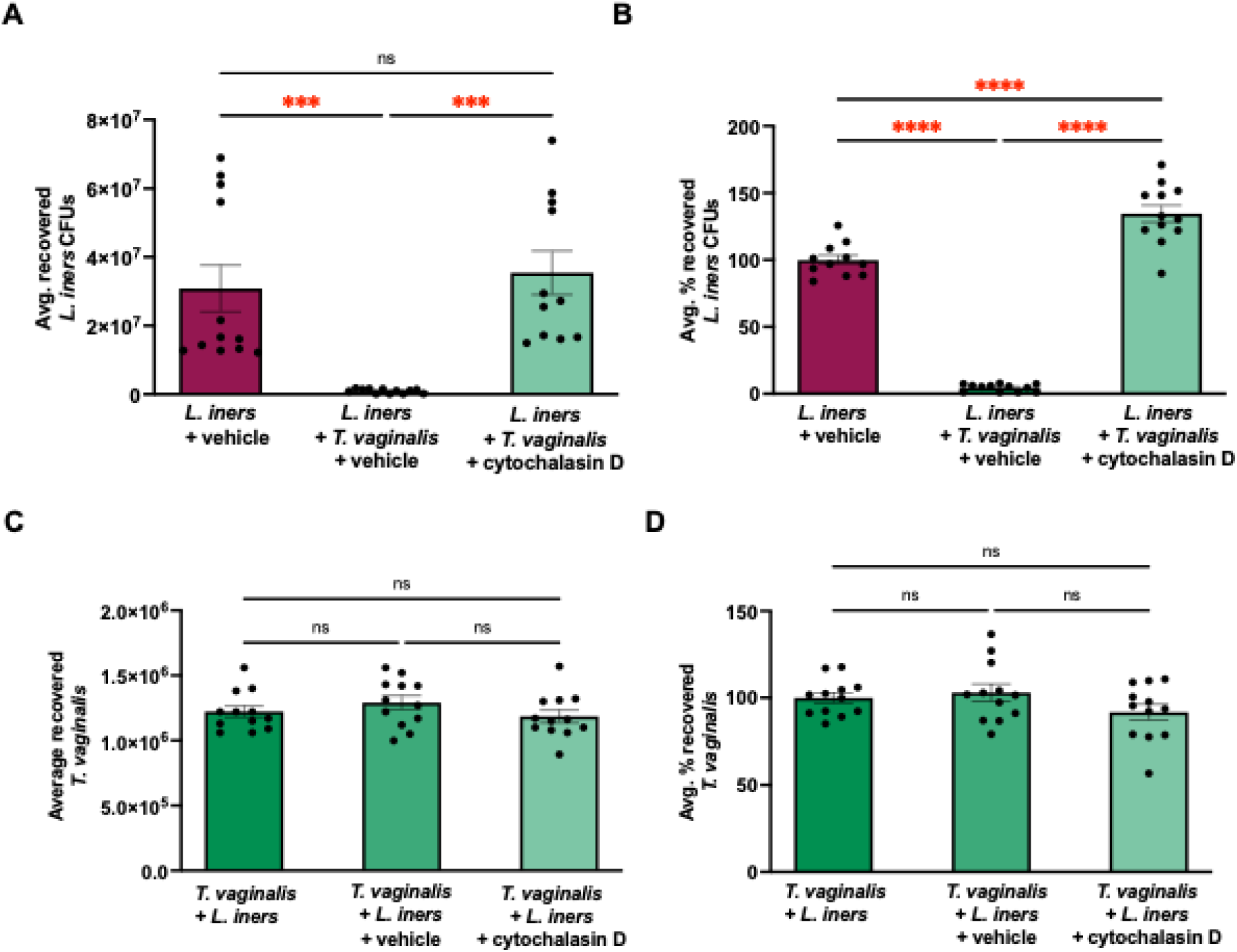
Cytochalasin D leads to a protective effect on *L. iners* viability. Co-incubation of *L. iners* with *T. vaginalis* was performed at MOIs of 9-43 for 3 hours under anaerobic conditions, in the presence of DMSO vehicle control or 60 µM cytochalasin D. (**A**) *L. iners* viability was quantified by measurement of colony forming units (CFUs) on CBA+CQ plates. (**B**) Percent CFUs normalized to the *L. iners*-only condition. (**C**) *T. vaginalis* viability was assessed via hemocytometer cell counts. (**D**) Percent viability normalized to the *T. vaginalis*-only control. Graphs show the pooled mean from three biologically independent experiments with the standard error of the mean displayed as error bars. Individual technical replicates are shown as dots. **** = p < 0.0001, *** = p < 0.001 ** = p < 0.01, * = p < 0.05, ns = not significant (p > 0.05).

## Discussion

Consistent with a prior study, we confirm that *L. iners* requires cysteine supplementation for robust growth in broth [32]. In this work, we expand on this finding by discovering that cysteine and glutamine supplementation of Columbia Blood agar plates dramatically improves detection of colony formation. Amino acid supplementation reduced the incubation time to quantify *L. iners* viability from over 72 hours to 48 hours. Using this improved colony formation assessment method to more accurately quantify *L. iners,* we were able to aliquot desired amounts of *L. iners* to reproducibly study its interaction with *T. vaginalis*. We first investigated early interactions between *T. vaginalis* and *L. iners* using ImageStream to visualize and quantify microbe-microbe interactions. Our analysis revealed that *T. vaginalis* binds to *L. iners* by our first time point of 30 minutes and that these interactions increased by 1 hour of co-incubation (**Fig. 1**).

To further visualize *T. vaginalis*-*L. iners* interactions, we employed scanning electron microscopy (SEM). These images captured a myriad of dynamic interactions between *T. vaginalis* and *L. iners* (**Fig. 2–4**). We identified the presence of membrane extensions that protruded from *T. vaginalis* towards *L. iners* (**Fig. 2A-2B** and **Fig. 2C-2D**). We also observed that these membrane protrusions formed lattice-like structures that appear to ensnare the bacteria, suggesting that *T. vaginalis* may actively recognize and respond to contact with *L. iners.* More dramatic reorganization of the *T. vaginalis* membrane in the process of beginning to engulf *L. iners* (**Fig. 2E-2F**) as well as after uptake of *L. iners* was captured (**Fig. 3**). These results are suggestive of phagocytosis-like behavior by *T. vaginalis* towards *L. iners.* Phagocytosis of whole yeast cells and other bacteria by *T. vaginalis* has been previously reported [37, 38], however, to our knowledge this is the first high resolution imaging of *T. vaginalis* in contact with *L. iners*. The changes observed in the *T. vaginalis* plasma membrane suggest a multi-stage interaction process that may include adherence, tethering, and subsequent uptake of the bacteria.

SEM analysis of *L. iners* morphology during co-incubation with *T. vaginalis* also revealed alterations to the *L. iners* surface (**Fig. 4**). Specifically, we observed a statistically significant increase in the percent of bacteria displaying a distinctive “zebra-like” striated pattern on the *L. iners* cell surface upon co-incubation with *T. vaginalis* compared to the *L. iners-*only condition (**Fig. 4B**). The *L. iners* striated membrane morphology could be indicative of increased cellular damage to *L. iners* caused by *T. vaginalis*. To further probe how this membrane alteration was inflicted, we utilized Cerillo Co-Culture Duets to allow the exchange of soluble factors between both microbes placed in adjacent chambers but provided physical separation between them. Using this system, we found that preventing *T. vaginalis* from binding to *L. iners* abolished the increased induction of the striated “zebra-like” pattern on the *L. iners* surface (**Fig. 5**). While the underlying cause of this bacterial phenotype remains to be elucidated, it may reflect mechanical damage arising from cycles of *T. vaginalis* binding and detachment to *L. iners* or membrane damage inflicted upon parasite attempts to engulf it. Alternatively, *L. iners* may be injured while outside of the parasite by exposure to proteases or potential saposin-like pore-forming proteins we have found to be released by *T. vaginalis* [39]. For example, saposin-like proteins are secreted by the eukaryotic pathogen *Entamoeba histolytica* and display antibacterial effects [40]. Diaz et al. characterized a specific *T. vaginalis* saposin-like family member, TvSaplip12, and found it had antibacterial activity against *Escherichia coli and Staphylococcus aureus* [41]. Additionally, Pinheiro et al. and Barnett et al. have found that *T. vaginalis* NlpC/P60 peptidoglycan hydrolases reduce the viability of *E. coli* and *Lactobacillus gasseri*, respectively [24, 42]. It is thus of interest to test the contribution of these enzymes towards *T. vaginalis* antibacterial activity against *L. iners*. Future studies are also needed to determine whether the zebra-like morphology we increasingly detected upon *L. iners* exposure to *T. vaginalis* is indicative of attempts of the bacteria to repair its membrane or if this phenotype represents a snapshot of early stages in *L. iners* lysis or metabolic dysfunction. Our findings indicate that *T. vaginalis* can directly damage *L. iners*, potentially through contact-dependent mechanisms or secreted effectors that may need to act in close range after *T. vaginalis* association with *L. iners*.

After identifying that *T. vaginalis* intimately adhered to *L. iners* and induced ultrastructural damage to the bacteria, we assessed the ultimate impact of *T. vaginalis*-*L. iners* interactions on the viability of both microbes. Total killing assays revealed that *T. vaginalis* actively kills *L. iners* (**Fig. 6A-6B**). *L. iners* viability was potently affected by co-incubation with *T. vaginalis*, with a 95% statistically significant decrease in recoverable CFUs compared to *L. iners* cultured alone (**Fig. 6B**). Interestingly, *T. vaginalis* viability in the *T. vaginalis* only control was similar to the amounts of the parasite recovered after exposure to *L. iners* (**Fig. 6B**). This is a significant finding, as the microbiological properties of *L. iners* may differ to those of other *Lactobacillus* species. For example, *L. crispatus* has been found to reduce the viability of the eukaryotic fungal pathogen *Candida albicans* [43, 44], whereas *L. iners* did not affect *T. vaginalis* viability. In contrast, the decrease in recoverable *L. iners* CFUs upon co-incubation with the parasite is consistent with the potential signs of bacterial entrapment, bacterial damage, and bacterial uptake observed in the SEM micrographs (**Fig. 2–5**). Together, these findings support a model in which *T. vaginalis* engages in an antagonistic interaction with *L. iners* and directly reduces its viability.

We also found that *T. vaginalis* potently killed *G. vaginalis*, leading to an 89% decrease in recovered *G. vaginalis* after co-incubation (**Fig. 6C**), whereas *P. bivia* viability was unaffected by the presence of *T. vaginalis* (**Fig. 6E**). Like our findings with *L. iners*, *T. vaginalis* viability was also unaffected by co-incubation with *G. vaginalis* and *P. bivia* (**Fig. 6D** and **Fig. 6F**). The capacity of *T. vaginalis* to reduce the amounts of *L. iners*, could have important implications for women that contract a *T. vaginalis* infection, wherein the parasite may shift the vaginal ecosystem toward a more dysbiotic state, as *L. iners* dominated cervicovaginal microbiomes are already increasingly prone to shift towards a bacterial vaginosis-like community [33, 34].

Since *T. vaginalis* can kill *G. vaginalis* but it does not exert antibacterial activity against *P. bivia*, two bacterial species increasingly detected during dysbiosis/bacterial vaginosis, it is likely that multiple cues and molecular factors govern the ability of *T. vaginalis* to coexist with certain cervicovaginal bacteria vs. trigger its antibacterial effects. These molecular cues may also change in differing host microenvironments. For example, although we observed loss of *G. vaginalis* viability upon *T. vaginalis*-*G. vaginalis* interactions, the study by Barnett et. al did not detect a loss of viability upon *G. vaginalis* exposure to *T. vaginalis* [45]. Our study utilized a 50/50 media mixture to give both microbes their optimal growth conditions whereas in the Barnett et al. study serum-free keratinocyte media (K-SFM) was utilized to mimic host cell conditions. Together, these results highlight that the use of microbiological vs. host media may contribute to differential antibacterial activity by *T. vaginalis*. Future comparative analysis under varying metabolic conditions and with different cervicovaginal bacteria could also help to pinpoint what makes *L. iners* particularly susceptible to *T. vaginalis*. The fact that *T. vaginalis* viability was not reduced by the presence of various cervicovaginal bacteria suggests a degree of parasite resilience or adaptation to co-inhabit the vaginal niche. Furthermore, the asymmetrical microbe-microbe interactions observed with *T. vaginalis*-*L. iners* and *T. vaginalis-G. vaginalis* may confer ecological advantages to *T. vaginalis*, allowing it to outcompete these bacteria when needed.

The dramatic appearance of thin-membrane protrusions extending from *T. vaginalis* towards *L. iners* and (**Fig. 2A-2B** and **Fig. 2C-2D**) and engulfment of *L. iners* (**Fig. 2C** and **Fig. 3**), suggested that membrane remodeling events driven by actin polymerization may contribute to *T. vaginalis* ability to kill *L. iners* (**Fig. 6**). We found that *T. vaginalis* antibacterial activity against *L. iners* was indeed dependent on actin cytoskeleton dynamics as treatment with the actin polymerization inhibitor cytochalasin D caused a complete protection of the bacteria (**Fig. 7A**). This is consistent with a prior study in which Pereira-Neves and Benchimol showed that treatment with cytochalasin D inhibited the uptake of yeast cells by *T. vaginalis*, highlighting the role of actin filaments in this process [37]. Importantly, cytochalasin D treatment did not significantly impact *T. vaginalis* viability under the conditions used (**Fig. 7B**), thus the loss of *T. vaginalis* antibacterial effects against *L. iners* does not arise as a result of nonspecific toxicity on the parasite, also consistent with [37].

We identified that *T. vaginalis* relies on an actin-dependent, contact-mediated mechanism to interact with and damage *L. iners*, possibly in part through phagocytosis. This adds mechanistic insight as to how the morphological changes we observed in both microbes could be mediated. For example, *T. vaginalis* membrane protrusions potentially arise as result of sensing *L. iners*. Some of the *T. vaginalis* membrane protrusions that we detected resemble the cytoneme-like membrane structures reported by Salas et al. upon *T. vaginalis*-*T. vaginalis* interactions [46]. However, we also observed more intricate crisscrossing of these membrane protrusions, in honeycomb-like structures that appear to help *T. vaginalis* capture the bacteria (**Fig. 2A-2B**) which to our knowledge is a novel observation. Additionally, the areas in which *L. iners* appears to be taken up by the parasite (**Fig. 3**) also have circular membrane rings resembling phagocytic cups. These mechanistic findings contribute to a growing understanding of how *T. vaginalis* interacts with vaginal microbiota and suggests that disrupting parasite cytoskeletal function could be a potential strategy to preserve beneficial bacterial populations such as that of *L. iners*, if further studies continue to uncover a protective role for it.

The ability of *T. vaginalis* to kill *L. iners*, a dominant member of the cervicovaginal microbiome has important implications for microbial competition, host immune modulation, and the pathogenesis of trichomoniasis. *L. iners* has been associated with both healthy and dysbiotic vaginal environments, and its removal or suppression by *T. vaginalis* may alter the cervicovaginal microbiome in ways that modify the host microenvironment to promote parasite infection or susceptibility to other pathogens, the later of which may also benefit *T. vaginalis* colonization. Additional microbe-microbe studies characterizing this synergy are needed to further explore this potential outcome. This study provides the first evidence that *T. vaginalis* directly interacts with and kills *L. iners* through an actin-dependent, contact-mediated mechanism. It is important to consider that while nitroimidazole drugs used to treat *T. vaginalis* infections are mostly effective at eliminating the parasite, the potential collateral disruption to the cervicovaginal microbiome by *T. vaginalis* infection*s*–such as perturbation to the amounts or depletion of *L. iners*–may persist beyond treatment. These cervicovaginal microbiome alterations could have lasting impacts on female health. Thus, in addition to the need of developing antiparasitic drugs against *T. vaginalis* in the face of drug resistance, design of therapeutics to disarm *T. vaginalis* antibacterial activity may also be necessary to preserve or restore microbial homeostasis after *T. vaginalis* colonization.

## Acknowledgements

This study was supported by a Research Project Grant to A.M.R from the National Institute on Minority Health and Health Disparities of the National Institutes of Health under Award Numbers S21MD010690 (SDSU HealthLINK Endowment) and U54MD012397 (SDSU HealthLINK Center). The content is solely the responsibility of the authors and does not necessarily represent the official views of the National Institutes of Health. This study was also supported by a Prebys Research Heroes Grant from the Prebys Foundation to A.M.R.

